# Acute Stress in Female Adolescent Rats Increases Anxiety-like but not Depression-like Behaviors

**DOI:** 10.1101/2025.11.20.689244

**Authors:** Jinglin Xiong, Ioanna Giannakou, Madison O’Bryan, Christin Hartman, Nancy Rempel-Clower

**Author notes:** Direct correspondence to: Nancy Rempel-Clower, Ph.D., Department of Psychology, Grinnell College, Grinnell, IA 50112. Shared first authorship (Jinglin Xiong, Ioanna Giannakou, Madison O’Bryan, Christin Hartman).

## Abstract

Adolescence is a critical developmental period where heightened stress reactivity and responsiveness increase susceptibility to psychiatric conditions like anxiety and depression. Studies with rodent models have primarily tested the effects of chronic, rather than acute, stress on the development of anxiety- and depression-like behaviors, and many have not included female subjects. The current study examined whether a single acute stress exposure in female rats can induce anxiety-like and depression-like behaviors, and whether the timing of the stress, early (postnatal day 28) versus mid-adolescence (postnatal day 36), influences these outcomes. We used a PTSD-relevant stress paradigm that paired acute immobilization with fox-urine predator odor. Female Sprague-Dawley rats completed a battery of behavioral assays to characterize anxiety-like (elevated plus maze, open field test) and depression-like behaviors (sucrose preference test, forced swim test) between 12-16 days post exposure. Acute stress in either early or mid-adolescence caused an increase in anxiety-like behavior on the elevated plus maze 15 days post-exposure. No increase in depression-like behaviors was observed at either age. Our findings suggest that a single exposure to a predator odor when paired with immobilization is sufficient to increase anxiety-like but not depression-like behaviors in female adolescent rats. These results provide evidence that acute stress during either early or mid-adolescence can lead to later anxiety in females.

## Introduction

Adolescence is a sensitive developmental window characterized by important physiological and behavioral changes (Cohen et al., 2013). Experiencing stress during this period can cause long-term neuroplastic changes resulting in emotional and behavioral disturbances (Gunnar, 2021; Perica & Luna, 2023; Romeo, 2010). Psychiatric disorders that emerge around late adolescence or early adulthood are heavily associated with stress experienced during this critical period, where the severity of the pathology can depend on the developmental timing of childhood or early adolescent trauma (Dunn et al., 2019). Moreover, females are disproportionately affected by psychiatric mood disorders (Remes et al., 2016) such that adolescent girls have higher rates of depression and anxiety than their male counterparts (Angold et al., 1999; Breslau et al., 2017). This pattern generalizes to non-human species: rodent studies consistently show that the impact of adolescent stress on anxiety- and depression-like behaviors depends on the sex of the animals For example, chronic stress exposure during adolescence induces more pronounced depression-like behaviors in females than males, with effects observed during late adolescence and adulthood (Bourke & Neigh, 2011; Ding et al., 2024). The heightened susceptibility to affective disorders in females highlights the importance of understanding the effects of stress during adolescence in a sex-specific manner.

Because chronic paradigms involve repeated exposures across weeks, they make it difficult to pinpoint when within adolescence is the most vulnerable to stress exposure. The precise timing of stress exposure during the adolescent period, particularly acute stress, was found to affect anxiety-like behaviors and stress hormonal response in rats (Cymerblit-Sabba et al., 2015) and may interact with known sex-specific effects. Adolescence in rats can be divided into three age intervals: early adolescence (weaning at postnatal day 21 (P21) to P34), mid-adolescence (onset of puberty, P34–46), and late adolescence (young adult, P46–59) (Tirelli et al., 2003). Research investigating the precise timing of acute stress exposure that may lead to psychiatric disorders during mid-to-late adolescence is limited, particularly in females (McCormick, 2022).

The current study aims to address this gap in the literature by introducing an acute-traumatic stressor at either early (P28) or mid (P36) adolescence and quantifying subsequent differences in anxiety- and depression-like behaviors during mid-to-late adolescence. We use a rodent model of post-traumatic stress disorder (PTSD) induced by simultaneous exposure to two evolutionarily relevant acute stressors (immobilization and presentation of an olfactory stimulus resembling a predator (Zoladz & Diamond, 2016)) to manipulate the duration of stress exposure more precisely and reveal which specific stages of adolescence might be the most vulnerable to the development of psychiatric disorders (Verbitsky et al., 2020). This combinatorial acute stress exposure leads to behavioral outcomes commonly seen in chronic stress models, such as increased anxiety-like behaviors and higher corticosterone levels, suggesting that this model is successful at modeling PTSD in rodents (Breton et al., 2021; Malin et al., 2023).

## Methods

### Subjects

Adolescent female Sprague-Dawley rats (Envigo, Indianapolis, IN) were tested in this study. Rats were matched by birthdate and pair-housed in a plastic cage with *ab libitum* access to food and water, on a 12/12-hour light/dark cycle (lights on at 07:00). Rats were randomly assigned to stress or control conditions by cage. Stress exposure, weight measurements, and behavioral tests were performed during the light period. All animal protocols were approved by the Grinnell College Institutional Animal Care and Use Committee and were conducted in accordance with published guidelines.

### Stress protocol

Rats assigned to the stress condition were exposed to an acute combinatory stressor one day after arrival at the facility, when they were either at postnatal day 28 (P28; early adolescence) or postnatal day 36 (P36; mid-adolescence) (**Figure 1**). Animals were removed from the housing room and kept with their cagemates in a clean cage free of bedding. Each rat was immobilized in a DecapiCone bag (Braintree Scientific, Inc., Braintree, MA) and placed next to its cagemate with their noses approximately 1 inch away from a cotton ball infused with 1mL of fox urine (Maine Outdoor Solutions, LLC, Hermon, Maine). The combined stressor of immobilization and predator odor lasted for 3 hours, from 9:00 to 12:00. Immediately following stress exposure, rats were placed in clean cages with their cagemates and allowed to self-groom. For three days following stress exposure, stress-exposed animals were kept in a separate housing room to minimize the transfer of stress to control animals. Rats in the control condition were weighed and handled according to the same schedule as those in the stress condition but otherwise remained in their home cages while rats in the stress condition were exposed to the stressor.

**Figure 1.**
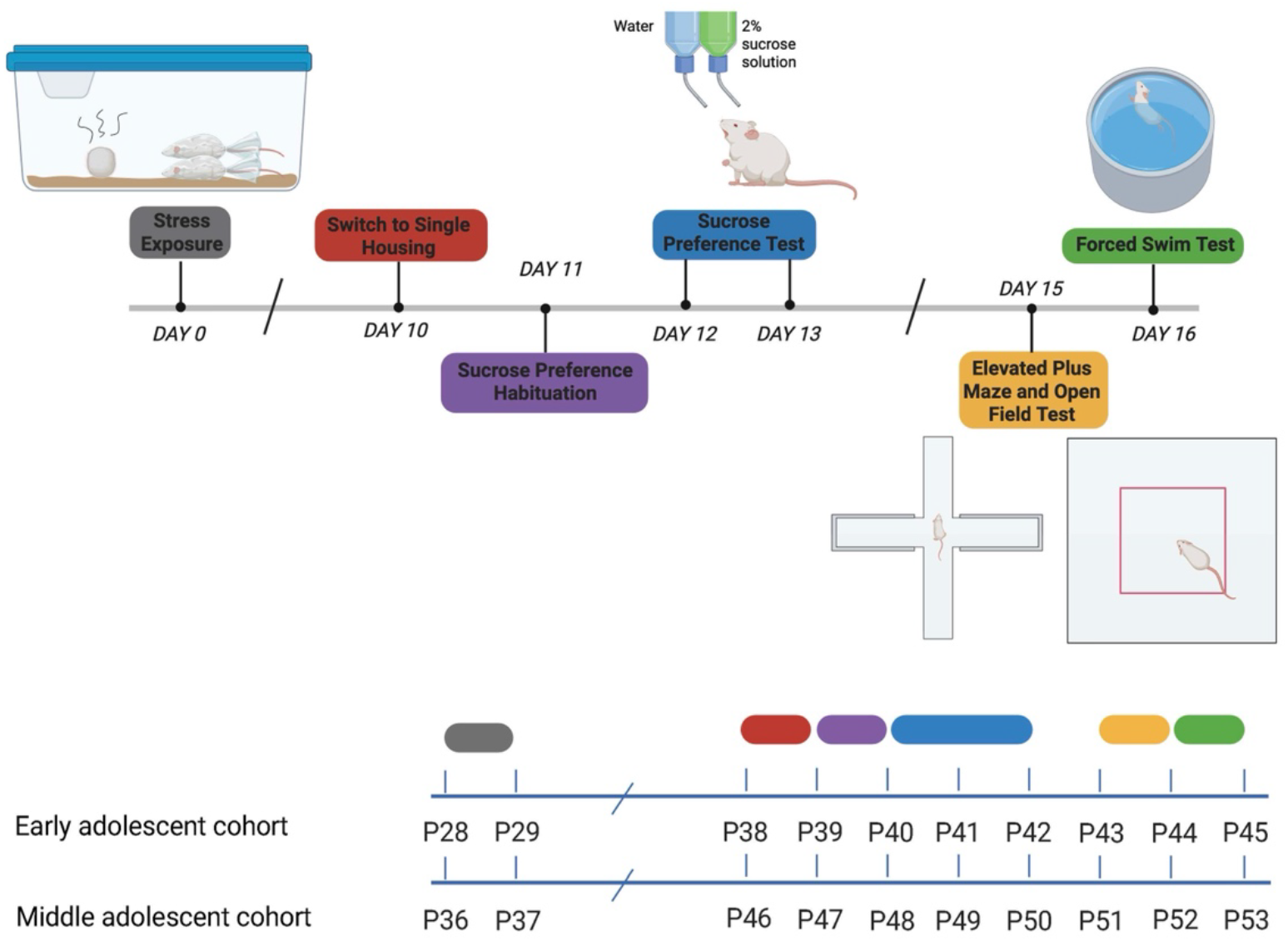
Experimental timeline of stress exposure and behavioral testing. Rats were subjected to a 3-h stress protocol on Day 0, after which they underwent sucrose preference habituation on Days 11 and sucrose preference test on Days 12–13. Anxiety-like behavior was then assessed in the elevated plus maze and open field test on Day 15, and depressive-like behavior was evaluated in the forced swim test on Day 16. Apparatus illustrations and timeline graphics were created with BioRender.

### Weighing and handling

Immediately before the stress exposure, each rat was weighed. Weight was recorded at the same time every day for a week following stress exposure. In addition to handling associated with weighing, rats were handled for 2 min 1 day prior to stress exposure, 1 day prior to the start of the sucrose preference habituation and testing, and 1 day prior to the elevated plus maze and open field task.

### Behavioral assays

Two age cohorts were run and tested at different developmental stages. Ten days after stress exposure, rats were switched to single housing and remained single-housed for all subsequent procedures. Behavioral testing was run in the order shown in **Figure 1**. Behavioral testing for the early-adolescent cohort occurred during mid adolescence, whereas testing for the mid-adolescent cohort occurred during late adolescence.

#### Sucrose Preference Test (SPT)

24 hours before the start of a habituation period, rats were simultaneously offered two water bottles, one with regular water, and one with a 2% sucrose solution for a 24-hour period. During this period, the position of the bottles was switched after 12 hours to control for side preference. After habituation, animals underwent a 48-hour testing period, with the position of the bottles switched after 24 hours. The weight of each bottle was recorded at the beginning and the end of the testing period to measure liquid consumption. Sucrose preference, a measure of anhedonia (Liu et al., 2018), was calculated as the percentage of sucrose consumption over the overall liquid consumption.

#### Elevated Plus Maze (EPM)

Rats were tested on the elevated plus maze in an isolated, quiet, and dimly illuminated room. The maze consisted of two open and two closed arms (50 cm x 10 cm). Each animal was habituated in the room for 10 minutes, followed by 5 minutes of exploration on the EPM. At the beginning of the test, the animal was placed in the center facing one of the open arms. Time spent in open arms and velocity were coded using the Noldus Ethovision XT 11 software. The number of entries into the open arms and head dips from open arms were manually coded by video analysis. Anxiety-like behavior was defined as a decrease in open-arm activities, including number of entries, total duration of time spent in the open arms, and number of head dips (Walf & Frye, 2007). Anxiety Index, a measure that integrates open-arm time and entries described by Rao et al. (2012), was calculated using the following equation:

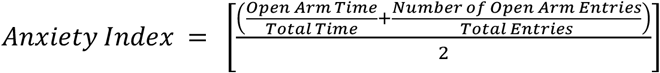

#### Open Field Test (OFT)

The OFT arena was an 82 cm x 82 cm box with 18 cm high walls. Each rat was habituated in the room for 10 minutes and then placed in the center of the apparatus and allowed to explore the open field for 5 minutes. All trials were run between 08:30 and 13:00 after each animal had finished the EPM. The center zone was defined as the inner 72 cm x 72 cm square. Time spent in the center zone, a reflection of anxiolytic behavior (Prut & Belzung, 2003), and velocity were coded using Noldus EthoVision XT 11.

#### Forced Swim Test (FST)

Twenty-four hours after an initial 12-minute exposure to the water, rats were tested in a 5-minute forced swim. During both the pretest and the test, rats were placed individually in a water-filled cylindrical plastic bucket measuring approximately 40 cm high and 30 cm in diameter. The depth of the water was 7-10 cm below the top rim of the bucket and the temperature was 25°C ± 1°C. Following both swims, the rat was removed from the water and dried with a towel before being returned to its home cage. Overhead cameras recorded video of the behavior, which was then coded manually by two researchers separately (96% interrater reliability) using a time sampling method (Detke et al., 1995). At the end of every 5-second period, behavior was coded at that moment as immobilization, climbing, or swimming. Increased levels of depression-like behavior are indicated by an increase in counts spent immobile, a decrease in counts spent climbing and swimming, and a decrease in latency to immobilization (Porsolt et al., 1978). We defined immobilization as minimal movement required only to keep the rat’s head above water, with no more than one leg moving (Gregus et al., 2005; Hong et al., 2012). Climbing was defined as rapid movement in which the animal’s body appears to be vertical, with its two arms moving against the wall (López-Rubalcava & Lucki, 2000; Detke et al., 1995). Finally, swimming was defined as regular movement with two or more feet in the water (Hong et al., 2012). We also measured latency to immobilization as the time until the rat was immobile for more than 3 sequential time samples or “counts”. For latency to immobility, scores were selected by only one scorer per subject at random.

### Statistical Analyses

All values are reported as mean ± SEM. Data for all behavioral measures were evaluated using SPSS for extreme outliers, defined as data points more than three box lengths away from the center of the data. In each behavioral task, any animal with a single measurement considered an extreme outlier was excluded from that measurement, but not other measurements in the task. Two-way ANOVAs were used to evaluate the effect of age and condition on animal performance on behavioral tasks, followed by post-hoc pairwise comparisons using Bonferroni’s correction method if the main effect or interaction was significant. For analysis of the effect of acute stress on body weight gain, a three-way mixed ANOVA was used, followed by simple main effect analysis. All statistical analyses were conducted using RStudio or SPSS. Graphs were created using Prism. Complete raw data files used for statistical analyses are available at OSF (DOI: 10.17605/OSF.IO/W7YD5). Full statistical results are provided in **Table S1**.

## Results

### Acute stress leads to less percentage gain in body weight

Body weight was measured daily for a week after the treatment to test the effectiveness of the stress protocol (**Figure 2**). A three-way mixed ANOVA yielded the main effects of stress condition (*F*(1,44) =21.52, *p* < 0.001), age (*F*(1, 44) = 582.44, *p* < 0.001), and the number of days after stress (*F*(3.13, 137.64) = 1558.35, *p* < 0.001). The main effect of the condition indicated that regardless of age, subjects in the acute stress condition had a lower percentage of body weight gain than control subjects. A significant interaction between age and the number of days after stress was observed (*F*(3.13, 137.64) = 217.92, *p* <0.001). No other significant interaction was found between condition, age, and the number of days post-stress.

**Figure 2.**
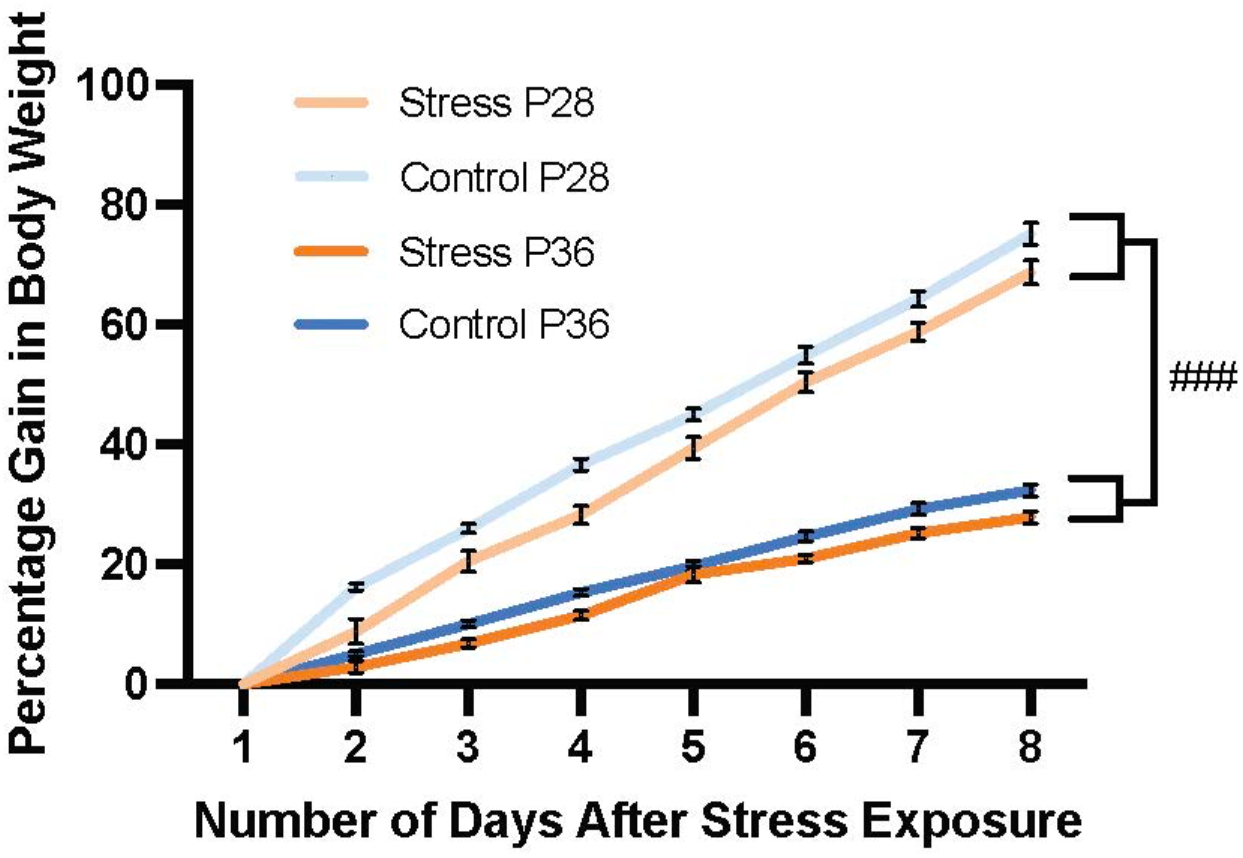
Percentage of weight gain from the day of stress exposure. Acute immobilization combined with predator odor exposure in early (P28) and mid-adolescent (P36) female rats reduced their percentage of body weight gain. ### *p* < 0.001 vs. control group regardless of age and the number of days after stress exposure. P28: control *n* = 12; stress *n* = 12. P36: control *n* = 12; stress *n* = 12. Error bars represent standard errors.

### Acute stress increases anxiety-like behaviors

Fifteen days after stress exposure, the rats completed two tasks to measure anxiety-like behavior: elevated plus maze and open field task. Rats were tested in the open field task 15 minutes after the elevated plus maze. In the elevated plus maze, acute stress led to an increase in anxiety-like behaviors and a decrease in explorative behaviors regardless of the age of stress exposure (**Figure 3**). Multiple two-way ANOVAs revealed that regardless of age, female rats exposed to acute stress spent less time in open arms (*F*(1, 43) = 9.76, *p* = 0.003) (**Figure 3A**), had fewer open arm entries (*F*(1, 43) = 7.06, *p* = 0.013) (**Figure 3B**), had fewer head dips on the open arms (*F*(1, 43) = 6.23, *p* = 0.011) (**Figure 3C**), and had higher anxiety index (*F*(1, 43) = 9.23, *p* = 0.004) (**Figure 3D**) than rats in the control condition. Bonferroni-corrected pairwise tests showed that adolescents exposed to stress at both ages increased anxiety-like behaviors. Animals exposed to stress during early adolescence spent less time in the open arms than controls (P28 stress vs P28 control, *p* = 0.036), and a similar reduction was observed for animals exposed to stress during mid-adolescence (P36 stress vs P36 control, *p* = 0.029; **Figure 3A**). Animals exposed to stress during early adolescence made significantly fewer open-arm entries than controls (*p* = 0.034), whereas the reduction after stress during mid-adolescence did not reach significance (*p* = 0.124; **Figure 3B**). Similarly, stress during early adolescence reduced the number of head dips on the open arms (*p* = 0.043), while stress during mid-adolescence did not affect this measure (*p* = 0.143; **Figure 3C**). Animals exposed to stress during early adolescence showed elevated anxiety index scores relative to controls (*p* = 0.021), and no significant difference was observed for animals exposed to stress during mid-adolescence (*p* = 0.066; **Figure 3D**).

**Figure 3.**
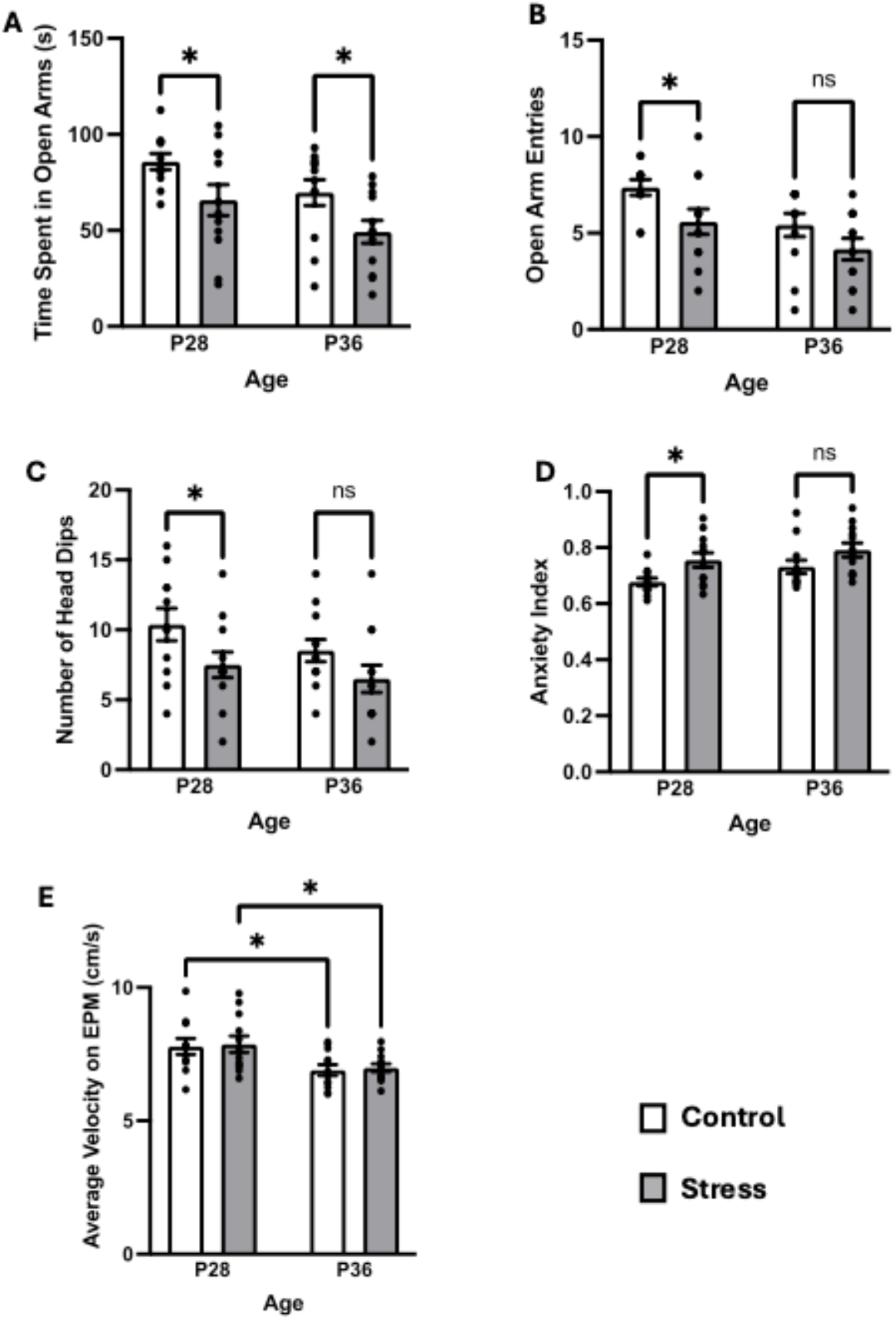
Anxiety-like behavior, explorative behavior, and general activity level on the elevated plus maze two weeks after stress exposure. (**A)** Time spent in open arms (s). **(B)** The number of open arm entries. **(C)** The number of head dips. **(D)** Anxiety index. **(E)** Average velocity in the elevated plus maze (cm/s). * indicates *p* <0.05. P28: control *n* = 11; stress *n* = 12. P36: control *n* = 12; stress *n* = 12. Error bars represent standard errors.

We also assessed the general activity level of rats in the elevated plus maze. Two-way ANOVAs yielded a significant main effect of age on average velocity (cm/s) in this task (*F*(1, 43) = 12.82, *p* <0.001) (**Figure 3E**). The average velocity was different by age in both conditions (P28 control vs P36 control, *p* = 0.016; P28 stress vs P36 stress, *p* = 0.014), indicating early adolescent rats moved at a faster speed than the mid-adolescent rats in the elevated plus maze. No effect of stress on average velocity was found in this task (*F*(1, 43) = 0.12, *p* = 0.727), suggesting that any anxiety-like changes we observed were not due to general activity level alterations.

In the open field task, no significant differences in anxiety-like behaviors between stress-exposed rats and control rats were detected. A two-way ANOVA yielded no main effect of condition on the time spent in the center zone (s) (*F*(1, 43) = 0.12, *p* = 0.729) (**Figure 4A**). We observed a significant main effect of age on velocity (*F*(1, 44) = 43.47, *p* < 0.001) (**Figure 4B**), with the early-adolescent animals moving at a faster speed than the mid-adolescent animals (P28 control vs P36 control, *p* < 0.001; P28 stress vs P36 stress, *p* < 0.001), consistent with their activity levels measured in the elevated plus maze.

**Figure 4.**
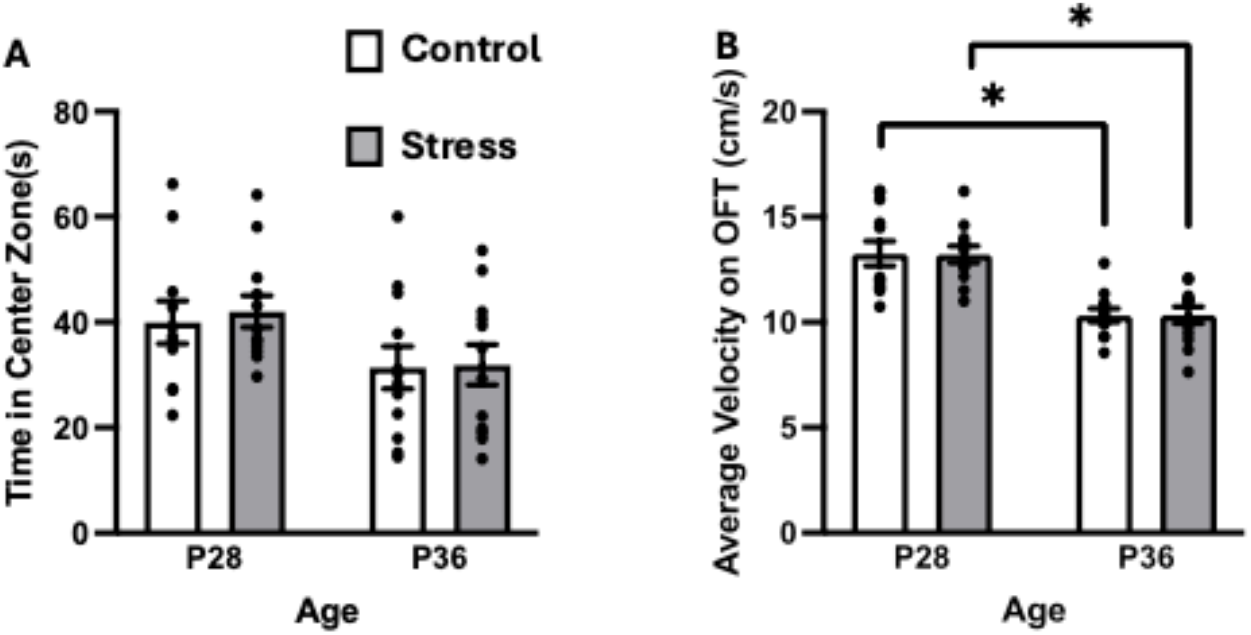
Anxiety-like behavior and general activity level on the open field test two weeks after stress exposure. **(A)** Time spent in the center zone (s), P28: control *n* = 11; stress *n* = 12. P36: control *n* = 12; stress *n* = 12. **(B)** Average velocity in the open field test (cm/s), P28: control *n* = 12; stress *n* = 12. P36: control *n* = 12; stress *n* = 12. * indicates *p* < 0.05. Error bars represent standard errors.

### Acute stress does not increase depression-like behavior

Ten days after the stress treatment, the rats were transitioned to single housing. Twenty-four hours later, they began a 24-hour habituation period in which they were presented with the water bottle and the sucrose bottle simultaneously. After that, they completed a 48-hour sucrose preference test. In the sucrose preference test, no differences in sucrose preference after acute stress exposure were observed between the two conditions (*F*(1, 37) = 0.08, *p* = 0.775) or the two age groups (*F*(1, 37) = 0.83, *p* = 0.368) (**Figure 5**). Multiple two-way ANOVAs yielded no significant interactions between age and condition for sucrose preference (*F*(1, 37) = 0.55, *p* = 0.463), total sucrose consumption (*F*(1, 44) < 0.001, *p* = 0.993), or total water consumption (*F*(1, 37) = 0.32, *p* = 0.577).

**Figure 5.**
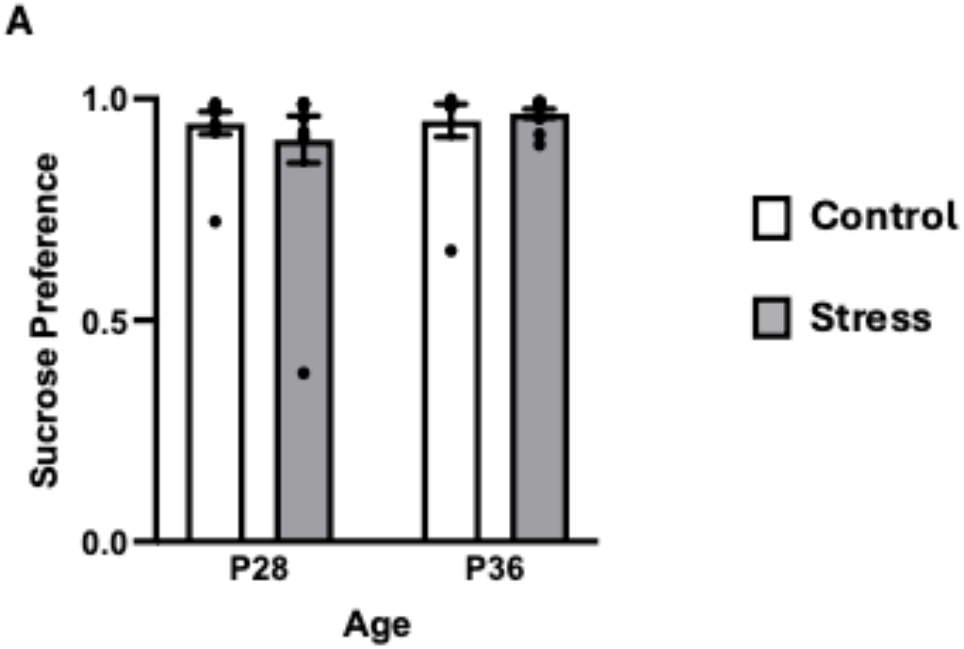
Depression-like behavior in the sucrose preference test two weeks after stress exposure. Percentage of sucrose consumption relative to overall liquid consumption in the sucrose preference test. P28: control *n* = 10; stress *n* = 11. P36: control *n* = 9; stress *n* = 11. Error bars represent standard errors.

Fifteen days after stress treatment, and after completing both anxiety-like behavioral tasks, rats completed a 12-minute pre-swim exposure, and a 5-minute forced swim test approximately 24 hours later. In the FST, a two-way ANOVA yielded a significant interaction between age and condition (*F*(1, 44) = 5.22, *p* = 0.027) for latency to immobility (**Figure 6A**), but no significant main effects of age (*F*(1, 44) = 0.492, *p* = 0.487) or condition (*F*(1, 44) = 0.35, *p* = 0.557). Bonferroni pairwise comparisons revealed that animals exposed to stress in mid-adolescence had significantly higher latency to immobility compared to controls (P36 control vs P36 stress, *p* = 0.048), as well as in comparison to animals exposed to stress in early adolescence (P28 stress vs P36 stress, *p* = 0.040). Multiple two-way ANOVAs yielded no significant interactions between age and condition for swimming (*F*(1,43) = 0.687, *p* = 0.412), climbing (*F*(1, 42) = 0.67, *p* = 0.416), or immobility (*F* (1, 43) = 2.27, *p* = 0.139) (**Figure 6B**). The significant interaction for latency to immobility suggests an age difference: animals exposed to stress during mid-adolescence appeared resilient by showing reduced depression-like behavior, whereas animals exposed to stress during early adolescence showed no change in depression-like behavior.

**Figure 6.**
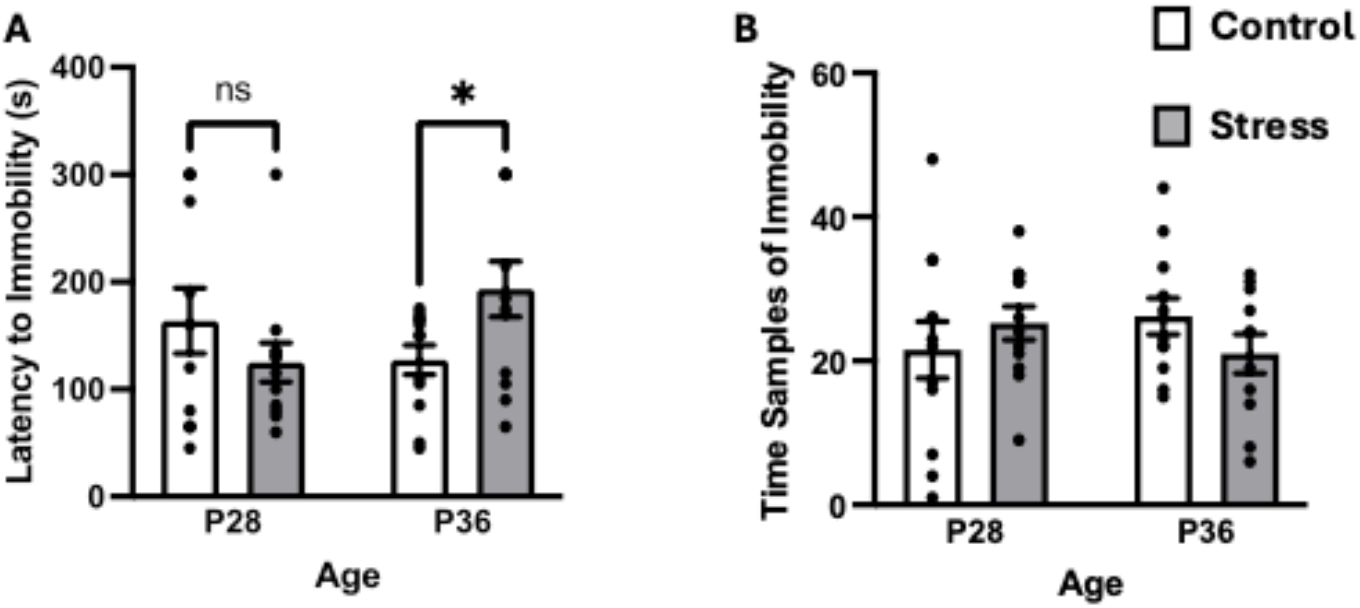
Depression-like behavior in the forced swim test two weeks after stress exposure. **(A)** Latency until the rat was immobile for more than three consecutive time samples in the forced swim test. P28: control *n* = 12; stress *n* = 12. P36: control *n* = 12; stress *n* = 12. **(B)** Total number of time samples scored as “immobile” in the forced swim test. P28: control *n* = 12; stress *n* = 12. P36: control *n* = 12; stress *n* = 11. * indicates *p* < 0.05. Error bars represent standard errors.

## Discussion

The goal of our study was to investigate the effects of acute stress in two distinct stages of adolescence in female rats on anxiety-like and depression-like behaviors. We found that a single exposure to a severe acute stressor in early (P28) or mid-adolescence (P36) was sufficient to elicit a robust physiological response. Additionally, animals exposed to acute stress during both early and mid-adolescence exhibited heightened anxiety-like behaviors two weeks later in mid-to-late adolescence, whereas this acute stress did not increase depression-like behaviors.

Previous research has shown that a combinatorial stress paradigm leads to reduced weight gain in both male and female juvenile rats after repeated stress exposure (Breton et al., 2021), as well as in adult male rats after a single exposure (Long et al., 2021). Our analysis revealed that female rats exposed to stress in both adolescent age groups gained significantly less weight compared to control animals. Reduced weight gains up to a week following a single stressful experience in adolescent females confirm that the stress exposure was effective. Although we did not directly measure the animals’ stress hormone levels, studies using similar stress paradigms (Breton et al., 2021; Long et al., 2021; Malin et al., 2023) have reported elevated plasma corticosterone levels in stressed animals for up to 200 minutes following exposure accompanying reduced weight gain in adolescents or weight loss in adults.

As predicted, acute stress exposure during adolescence caused an increase in anxiety-like behavior in our subjects, regardless of whether the stress occurred in early or mid-adolescence. Female rats exposed to a single acute immobilization stress in early adolescence exhibited robust anxiogenic effects when tested 15 days later on the elevated plus maze, including significant reductions in open-arm exploration and changes in multiple anxiety-related measures. Likewise, rats exposed to stress in mid-adolescence also showed increased anxiety-like behavior 15 days post-stress, as shown by a significant decrease in time spent in the open arms. Our findings indicate that a single acute stressor during either early or mid-adolescence is sufficient to produce long-term increases in anxiety-like behavior in female rats. These results are consistent with a broad literature showing that acute stressors during early adolescence have been shown to induce persistent anxiogenic effects in later life. For example, a single footshock delivered at P29 increased avoidance and anxiety-like behavior in adult male rats (Lovelock & Deak, 2019). Similarly, adolescent male rats exposed to acute restraint stress displayed elevated anxiety-like behaviors and corticosterone levels in adulthood (Sánchez-Marín et al., 2022). There is also evidence that stress in mid-adolescence can cause an increase in anxiety-related outcomes: male rats subjected to an acute restraint stress in mid-adolescence showed increased anxiety-like behavior when tested 10 days later, whereas those stressed in early adolescence were stress-resilient (Campos-Cardoso et al., 2023). Our results extend this literature by demonstrating that in female rats, a single acute stress in both early and mid-adolescence leads to heightened anxiety-like responses 15 days following exposure. This highlights both early and mid-adolescence as vulnerable developmental periods during which even brief stressors can impart lasting emotional consequences.

We found that exposure during adolescence to a single, highly traumatic event, immobilization in the presence of predator odor is sufficient to trigger reduced weight gain and changes in anxiety-like behaviors. This intense paired stressor is an effective model for studying PTSD (Zoladz & Diamond, 2016). However, not all adolescent short-term stressors can cause adverse emotional outcomes. When the stressor is moderate and predictable, it can be beneficial: a single 3-h of immobilization in a neutral context increased prosocial behavior via heightened oxytocin signaling in male rats (Muroy et al., 2016), and a short predictable stress for 12 days during adolescence had a protective effect on anxiety-like behaviors in both sexes (de Melo et al., 2020). Adverse emotional outcomes emerge when the stressors are severe enough. Previous research has shown that the combination of immobilization and predator odor stress paradigm in adult male and female rats can produce robust stress responses including increased anxiety-like behaviors and PTSD-related startle responses (Long et al., 2021; Malin et al., 2023). Consistent with our findings, even a single 15-min episode of simultaneous predator odor and restraint during adolescence induced persistent anxiety-like behavior in male rats (Borodovitsyna et al., 2018). Our study extends these findings to female adolescent rats, showing that one brief, high-intensity stressor is sufficient to cause both immediate and lasting maladaptive outcomes. However, we note that rats were switched to single housing 10 days after the stress exposure and remained single-housed throughout all behavioral testing. This switch from pair housing to single housing can be considered as social isolation, a potential second hit in a two-hit stress model. Prior work showed that stress combined with, or followed by, isolation can exacerbate behavioral consequences of the stressor (Sailer et al., 2022; Si et al., 2023). In our context, the increase in anxiety-like behaviors could reflect the interaction between the initial stress exposure and later social isolation, whereas the reduced weight gain, measured before isolation began, likely reflects the effects of acute stressor alone. Although isolation was introduced for logistical reasons, the sequence of a severe trauma followed by a period of social isolation may mirror aspects of human PTSD patients.

Our study suggests that a single exposure of adolescent female rats to a combinatory stressor is not sufficient to elicit depression-like behaviors. This observation differs from prior studies applying chronic stressors, reporting that repeated corticosterone injections in adult male animals increased learned helplessness (Gregus et al., 2005; Johnson et al., 2006) and that chronic restraint stress in adolescence elevated anhedonia in both sexes (Eiland et al., 2012). This variability in behavioral outcomes after exposure to stress might be impacted by stressor intensity and duration (Suvrathan et al., 2010), as different types of stressors often lead to distinct depression subtypes (Bai et al., 2014). Additionally, other studies suggest sex-specific resistance to depression-like behaviors, by reporting that chronic social stressors during adolescence resulted in elevated or sustained sucrose preference in adult female but not male rats (Hong et al., 2012; Renda et al., 2024). These sex differences may stem from stress-related effects on hormonal responses (Verma et al., 2010). In line with these observations, our results suggest that female adolescent rats may be resistant to developing depression-like behaviors after acute stress exposure. Our finding of higher latency to immobility during the forced swim test caused by stress during mid-adolescence further supports this hypothesis. While this finding is promising, it represents only a single behavioral measure. Future studies should use more robust behavioral measures and isolate the effects of sex, stressor duration, and developmental timeframe to better understand the mechanisms of resilience to development of depression-like behaviors.

The current study has certain limitations that should be addressed in future studies. First, the behavior of the two age groups was tested at two different chronological ages. To keep a consistent post-stress interval, rats exposed to stress in early adolescence were assessed during P40-44, whereas those in mid-adolescence were assessed at during P48-52. In contrast, Cymerblit-Sabba et al. (2015) exposed rats to acute stress at multiple developmental stages but tested every adolescent-stress cohort at the same adult age (P127). Although we do not anticipate large behavioral differences between P40 and P48, pubertal endocrine fluctuations and estrous cycle during this adolescent stage could affect anxiety-like behavior that we failed to capture (Pestana & Graham, 2024). Hormonal changes were not measured to avoid inducing additional stressors to the animals. Second, because behaviors were tested at one timepoint, we cannot confirm whether the observed increases in anxiety-like behaviors are persistent. Prior work found early adolescent stress causes increased anxiety-like behaviors in adult rats (Lovelock & Deak 2019; Sánchez-Marín et al. 2022), so we expect the increase in anxiety-like behaviors to persist into adulthood. Similarly, even though we did not observe depression-like behaviors at the time point of testing, it is possible that such behavioral changes might develop at later time points.

In summary, this study demonstrates that a single acute combinatory stress exposure in female adolescent rats increases anxiety-like, but not depression-like behavior measured later in adolescence. These changes are likely due to stress-induced neuronal alterations in brain regions involved in emotional processing (Kolb & Gibb, 2015). For example, acute stress exposure induces dendritic atrophy in both the hippocampus (Yang et al., 2020) and the medial prefrontal cortex (mPFC; Koskinen et al., 2020). In parallel, acute stress exposure increases glutamate release in the basolateral amygdala (BLA), driving HPA axis activation (Lynn & Maillé, 2017). This is further amplified in adolescents due to higher glucocorticoid levels after stress, potentially contributing to the increased anxiety-like behaviors seen in younger individuals (Romeo et al., 2013). Stress effects on neuronal architecture also differ by sex, with males showing dendritic restrictions in the hippocampus and the mPFC, and females displaying either no change or hypertrophy (Bowman et al., 2025). Therefore, future studies should explore neuronal changes in stress-related brain regions, while accounting for sex differences, to better understand the link between acute stress and behavioral outcomes in adolescent female rats.

## Conclusions

The current findings suggest nuances in the complex relationship between exposure to stress and the development of psychological disorders later in life. Our study identified both early adolescence and mid-adolescence as vulnerable periods for the emergence of stress-induced anxiety-like, but not depression-like behaviors in females. The lasting effects of a single PTSD-resembling stressor highlight the impact of both the intensity and the duration of the stress exposure in shaping the behavior of animals later in adolescence. The different patterns observed between anxiety and depression-like behaviors emphasize the importance of identifying distinct types of stressors that may differentially trigger the development of specific psychiatric disorders. These findings have important implications for human mental health, as adolescence is recognized as a sensitive period for the development of anxiety disorders, particularly among females (McLaughlin et al., 2013; Gerhard et al., 2021). Therefore, gaining insight into the consequences of exposure to stressors that occur during adolescence could guide the improvement of diagnostic tools for youth at heightened risk.

